# Molecular mechanisms of plastic biodegradation by the fungus *Clonostachys rosea*

**DOI:** 10.1101/2025.01.28.635403

**Authors:** Victor Gambarini, Nikolai Pavlov, Paul Young, Stephanie Dawes, Arnaud Auffret, Joanne M. Kingsbury, Lloyd A. Donaldson, Dawn A. Smith, Louise Weaver, Olga Pantos, Kim M. Handley, Gavin Lear

## Abstract

Microbial degradation provides an avenue for the remediation of select plastic polymers contributing to the urgent environmental problem of global plastic pollution. We demonstrate the degradation of polycaprolactone (PCL) by *Clonostachys rosea* and elucidate its underlying molecular mechanisms. We constructed the genome of this fungal strain and monitored changes in gene expression when exposed to PCL. Twelve genes linked to PCL degradation were found in the genome of *C. rosea*, and some of them were upregulated in the presence of the plastic, including genes coding for two cutinases. We heterologously expressed the enzymes coded by both genes and confirmed their activity against PCL polymers. We also demonstrate that one of the enzymes was active against polyethylene terephthalate (PET) polymers. Glucose inhibited the expression of both genes, completely halting the plastic biodegradation process, possibly serving as a preferred and readily metabolisable carbon source compared with PCL. We confirm the presence of key metabolic pathways linked to PCL degradation in *C. rosea*, including fatty acid degradation, providing further evidence of the mechanisms central to plastic biodegradation.

## INTRODUCTION

Plastic pollution is an urgent global environmental problem due to the extensive use and mismanagement of synthetic plastics and their persistence in the environment. Fish, seabirds, turtles, and marine mammals often mistake plastic debris for food, leading to ingestion [1–3], which causes gastrointestinal blockages or damages organs, for example, by plastic-induced fibrosis [4], in some cases resulting in starvation and death. Meanwhile, entanglement in plastic debris can cause strangulation, injuries, and hamper mobility, reducing survival rates [5, 6]. Plastic pollution also poses significant risks to human health. Plastics can release toxic chemical additives, such as phthalates, bisphenol A (BPA), and polychlorinated biphenyls (PCBs), which, if ingested, can cause endocrine disruption, reproductive problems, and have carcinogenic effects [7, 8].

Conventional management, such as recycling, remains insufficient to address the global plastic waste problem [9, 10]. Consequently, there is a growing interest in alternative plastic treatment methods, including biodegradation. Fungi have emerged as potential candidates for plastic degradation due to their diverse enzymatic capabilities, including the enzymatic degradation of recalcitrant organic substances [11–14]. Fungal enzyme classes, including laccases, peroxidases, esterases, lipases, proteases and ureases, have shown remarkable efficacy in degrading various plastics under laboratory conditions [13, 14], predominantly associated with members of the Basidiomycetes [15, 16] and Ascomycetes [17–19].

Fungal laccases [20, 21] and peroxidases [22] have shown some potential in degrading polyethylene (PE) and polyvinyl chloride (PVC), two of the most commonly used plastics, whereas esterases, such as cutinases and lipases, are effective in degrading polyethylene terephthalate [23], polycaprolactone (PCL) [24, 25], and polyurethane (PUR) [26]. Additionally, fungal proteases have shown potential for polylactic acid (PLA) degradation [27]. The diverse fungal enzymes involved in plastic degradation highlight fungi’s versatility and ability to target different types of plastics.

Despite the progress in understanding fungal enzymatic systems related to plastic degradation, identifying new microorganisms capable of efficiently degrading plastics, as well as the genetic machinery by which degradation is mediated, is of great interest. *Clonostachys rosea* (phylum Ascomycota), a saprophytic filamentous fungus, has emerged as a potential candidate for plastic degradation [28, 29]. Bainier [30] was the first to characterise *C. rosea*, previously known as *Gliocladium roseum*. Schroers*, et al.* [31] discovered that the morphology, ecology, teleomorph, and DNA sequences from the 28S rRNA gene of *G. roseum* were very distinct from other *Gliocladium* species, so reclassified *G. roseum* as *C. rosea*. It has been isolated from soil, plants (such as barley, onion, strawberry, rose and cocoa), insects, nematodes, and even wine [32–34]. This wide distribution and ability to colonise different environments highlight the ecological flexibility of *C. rosea* [35].

Understanding the biological characteristics of *C. rosea* is crucial for harnessing its potential applications, such as for biocontrol and biodegradation. Genomic studies provide insights into the biology of *C. rosea*. Its genome contains more than 14,000 annotated protein-coding genes [36, 37], including cell wall-degrading enzymes (CWDEs), polyketide synthases, ABC transporters, and monooxygenases [38]. Two studies have isolated and characterised *C. rosea* strains that can degrade polycaprolactone (PCL) and polyurethane (PU). Barratt*, et al.* [28] found that *C. rosea* was one of the key fungal strains responsible for the degradation of soil-buried polyester polyurethane. Urbanek*, et al.* [29] isolated *C. rosea* strain 16G from the Arctic, which exhibited the capacity to degrade 53% of PCL film within 30 days at 28 °C; a remarkable degradation efficiency even observed at lower temperatures (20 °C and 21 °C). These findings highlight the potential of *C. rosea* to be an effective degrader of plastic waste. Yet, the biological attributes of *C. rosea* that enable it to degrade plastic remain unclear.

We investigated the plastic-degrading capabilities of a strain of *C. rosea*. Firstly, we explored the capacity of this strain to degrade an extended range of plastics. After confirming the strain’s ability to degrade PCL, we explored the genetic and enzymatic systems that contribute to the PCL-degrading capabilities of *C. rosea* under various environmental conditions. The findings from this study provide insights into the genetic mechanisms, metabolic pathways, and enzymes used for fungi to degrade plastics, as well as the environmental variables that influence the plastic degradation process.

## MATERIALS AND METHODS

### Plastic polymers

PCL and PS were acquired from Sigma Aldrich (USA; products 440752 and 331651, respectively). Both PCL and PS were in pellet form and had average molecular weights of 14,000 and 35,000 g/mol, respectively. PLA pellets (Ingeo™ Biopolymer 3052D grade PLA) had an average molecular weight of 116,000 g/mol and were purchased from NatureWorks LLC, USA. Amorphous PET film (250 µM thick) was obtained from Goodfellow Cambridge Ltd (Huntingdon, UK) and cut into 2mm x 5 mm pieces. PET microparticles were prepared from this film by milling under liquid nitrogen in a SPEX Sample Prep 6970 EFM Freezer Mill at 10 cycles/second for a total grinding time of 25 minutes. The microparticles were fractionated using an Octagon 200CL Sieve Shaker (Endecotts Ltd, UK) through a series of sieves of descending size pores of 500 µm, 355 µm, 125 µm, and 63 µm (Endecotts Ltd, UK). Microparticles sized between 125 µm and 355 µm were used for assays.

### Preparation of polymer emulsions

PE liquid emulsions (Aquacer 1063) were a kind donation from ResChem (New Zealand). To prepare PCL, PS, and PLA polymer emulsions, 6 g of the respective polymer was dissolved in 100 mL dichloromethane (DCM) (Merck, Germany) at room temperature for 48 h. The solution was mixed with 300 mL of ddH_2_O and 1.2 mL of dishwashing liquid (Ecostore, New Zealand) and sonicated for 5 minutes at 50% amplitude, with one-second ON and OFF pulses, using a Q700 sonicator with a 1/2” diameter probe (Qsonica, Newtown, CT, USA). DCM was then evaporated by stirring at 50 °C in a fume hood for one hour. After DCM evaporation, the solution was cooled to room temperature.

### Genome characterisation

The strain of *C. rosea* utilised in our study was isolated in Rotorua, New Zealand, and provided by the Scion Research Institute in Rotorua, New Zealand. To isolate DNA from *C. rosea*, we used a PowerSoil® DNA Isolation Kit (MoBio Laboratories, Carlsbad, CA, USA). First, the fungal strain was grown on Potato Dextrose Agar (PDA) plates at 25°C for five days, and then approximately 200 mg of mycelium was scraped off the plate using a sterile loop and transferred to a 2 mL PowerBead tube. We followed the manufacturer’s instructions for DNA extraction, except for the mechanical lysis step, which was performed using a TissueLyzer II (Qiagen, Valencia, CA, USA) for 2 min at 30 Hz.

After extraction, the genomic DNA was cleaned and concentrated using a DNA Clean & Concentrator-5 kit (Zymo Research, Irvine, CA, USA), following the manufacturer’s instructions and the pellet was resuspended in UltraPure DNase/Rnase-Free Distilled Water (20 µl; Cat No. 10977015; Invitrogen, Thermo Fisher Scientific, Waltham, MA, USA). The quality and quantity of the extracted genomic DNA were initially assessed using a Qubit 2.0 Fluorometer (Invitrogen, Carlsbad, CA, USA) and agarose gel electrophoresis to ensure the DNA was of high integrity and of sufficient concentration for DNA sequencing. Genomic DNA was stored for downstream applications at –80 °C.

Whole genome sequencing was performed by Auckland Genomics Ltd. (University of Auckland, NZ). Auckland Genomics performed library preparation and sequenced the sample DNA on an Illumina MiSeq platform using a 500-cycle MiSeq Nano 2 x 250 bp kit, as per the manufacturer’s protocols.

### Biodegradation assays on plastic-emulsion agar plates

M9 minimal agar was prepared by adding 11.28 g of M9, Minimal Salts, 5X concentrate (Merck, New Zealand; product M6030) to 1 L of distilled water and autoclaving for 20 minutes at 121 °C. Following preparation, the medium was cooled to approximately 50°C. At this temperature, the previously prepared polymer emulsion was incorporated into the M9 medium at 50 mL of emulsion per litre of medium. The mixture was stirred thoroughly to ensure a uniform distribution of the polymer emulsion within the agar medium. Approximately 20 mL of this medium was dispensed into sterile 9 cm plastic petri dishes under aseptic conditions and allowed to solidify at room temperature.

*C. rosea* was inoculated onto the prepared agar plates’ surface by transferring a fungal mycelium plug using a sterile cork borer. The plates were incubated at room temperature (18-24 °C) and monitored daily, visually inspecting for microbial growth and any clear ‘halos’ in the plastic emulsion, indicative of plastic degradation. We did not test PET degradation using emulsions because PET was insoluble in all the solvents we evaluated.

### PCL biodegradation assays, incorporating varying carbon substrates

As other studies have found that adding supplementary carbon sources could improve the plastic degradation performance of some fungal species [39, 40], we tested the PCL degradation rates by *C. rosea* using different carbon sources. M9 minimal agar plates containing emulsified PCL were supplemented with either 1% (w/v) glucose, tryptic soya agar (TSA; Merck, Germany), citrate (Merck, Germany), potato starch (Merck, Germany), gelatine (Fisher Scientific, UK), fumarate (Merck, Germany), or cellulose (Sigma Aldrich, USA) to assess their potential influence on PCL biodegradation. Triplicate plates were inoculated with agar plugs of *C. rosea,* incubated at 25°C for 20 days, and monitored daily for signs of microbial growth and PCL degradation. Where observed, clearance zones were photographed to quantify the extent of biodegradation under the influence of each substrate. Images were processed using ImageJ software (NIH, USA), and the “measure” function was used to determine the diameter of the clear zones.

### RNA-Seq experiment

To identify the gene expression patterns associated with PCL degradation, we aimed to compare treatments in which the PCL degradation phenotype is either present or absent. Therefore, noting the inhibitory effect of glucose for PCL degradation by *C. rosea,* we cultured the organism under three media conditions: (i) PCL – M9 minimal agar supplemented with emulsified polycaprolactone (PCL); (ii) GLU – M9 minimal agar supplemented only with 1% (w/v) glucose (i.e., no PCL); (iii) GLUPCL – M9 minimal agar supplemented with both 1% (w/v) glucose and emulsified PCL. For each treatment (PCL, GLU, and GLUPCL), a plug of *C. rosea*, previously incubated under the same conditions, was transferred to Petri dishes to initiate the culture and incubated at 25 °C for six days. Each of the three treatments was replicated five times, yielding 15 samples. Six days post-inoculation, the fungus was collected from each Petri dish using a sterile scalpel and transferred into separate micro-centrifuge tubes. These were immediately flash-frozen in liquid nitrogen to preserve the integrity of the cellular RNA.

### RNA extraction

The preserved *C. rosea* RNA was extracted using an RNeasy Mini Kit (Qiagen, Hilden, Germany) according to the manufacturer’s instructions, with some modifications. Before starting, β-mercaptoethanol (β-ME) was added to Buffer RLT at a ratio of 10 μl β-ME per 1 ml Buffer RLT. Buffer RPE was prepared by adding four volumes of ethanol (96-100%) to obtain a working solution.

To extract RNA, 60 mg of flash-frozen fungal biomass was weighed in 2 mL microfuge tubes and then subjected to cell lysis. The cell lysis process involved using a pre-chilled stainless steel adapter block, 2.8 mm stainless steel grinding balls, and a Geno/Grinder tissue homogeniser (SPEX SamplePrep, Metuchen, NJ, USA). Cell lysis was performed at 1050 strokes/min for 1 min, followed by the addition of 700 μl Buffer RLT. The samples were ground twice at the same rate for 1 min each time and then ground once more for 2 min. After grinding, the samples were allowed to thaw at room temperature for 5 min. The RNA extraction process continued using a Qiagen RNeasy Mini Kit, following the manufacturer’s instructions, and eluting in RNAse-free water.

### RNA sequencing

Transcriptome sequencing was performed at Auckland Genomics, the University of Auckland, NZ. Before sequencing, the concentration and purity of the RNA were determined using a NanoDrop spectrophotometer (Thermo Fisher Scientific, Waltham, MA, USA), and the integrity of the RNA was assessed with an Agilent 2100 Bioanalyzer (Agilent Technologies, Santa Clara, CA, USA). Library preparation for sequencing was conducted using the Illumina Stranded mRNA Prep Kit according to the manufacturer’s instructions (Illumina, Inc., San Diego, CA, USA).

Quality control checks were performed on the prepared library using a Bioanalyzer 2100 to confirm appropriate fragment size distribution, and the library was quantified using a Qubit 2.0 Fluorometer (Invitrogen, Carlsbad, CA, USA). Normalization and pooling of the libraries for simultaneous sequencing were done as required by the sequencing facility, and the final pool underwent another round of quality control. The RNA library was then sequenced on an Illumina HiSeq platform utilising a 2×150 bp paired-end sequencing strategy.

### Film weight loss measurements

To confirm that the ability of *C. rosea* to degrade PCL is not limited to PCL emulsions, we fabricated PCL films by melting the polymer in a 70 °C water bath and compressing it with glass, resulting in rectangular films measuring approximately 10 x 20 mm and weighing close to 0.1 g. Films were sterilised by exposure to 70% ethanol for 30 min. A biodegradation experiment was conducted to measure the weight loss of the polymer after incubation with *C. rosea*, using three distinct liquid media conditions: 1) 11.28 g/L of M9; 2) 11.28 g/L of M9 and 4 g/L of yeast extract (yeast extract can frequently improve biodegradation rates [41]); 3) 11.28 g/L of M9 and 4 g/L of inhibitory glucose (and no yeast extract). For each treatment, four replicate and two control flasks were prepared (i.e., containing only the media, with no fungal inoculation). In each 600 mL Erlenmeyer flask, 100 mL of the specific medium was introduced. An actively growing plug of *C. rosea* mycelium was added to each flask (except for the control samples), incubated for 180 days at 28 °C, and agitated at 150 RPM.

Fungal mycelia were removed from the plastic films using sterile tweezers. The plastic films were then sterilised using a 70% ethanol solution for 10 minutes, followed by three rinses in sterile distilled water to remove residual ethanol and air-drying under a sterile laminar flow hood.

For Scanning Electron Microscopy (SEM) imaging, sections of the plastic films were cut and attached to aluminium stubs using adhesive carbon tabs, and sputter coated with 12 nm of chromium, before examination using a JEOL 6700F field emission SEM at an accelerating voltage of 3 kV before examination at 1000x magnification.

### Heterologous expression and protein purification

Two genes encoding the putative cutinases g9562 and g16887, upregulated in the presence of PCL, were synthesised and cloned into pET-28a(+)-TEV plasmids by GenScript Biotech PTE Ltd. (Singapore). The cutinases genes were cloned in the Ndel/EcoRI restriction enzyme sites of the plasmid, with the expressed protein containing an N-terminal hexa-histidine affinity tag and recombinant tobacco etch virus protease recognition site (rTEV). The plasmids (3 µL) were inserted via transformation into SHuffle T7 chemically competent *Escherichia coli* cells (New England Biolabs, Auckland, NZ) using a standard heat shock protocol. The transformed cells were plated on LB kanamycin plates and incubated for 24h at 30 °C. A single colony was cultured overnight in 10 mL of LB broth supplemented with kanamycin at 30°C and then transferred to 2 L baffled culture flasks containing 1 L of LB broth medium supplemented with kanamycin. The cultures were grown at 30°C with shaking to an OD_600nm_ of ∼0.8 before induction with 0.3 mM isopropyl-β-D-1-thiogalctopyranoside (IPTG). The cultures were transferred to 18 °C and incubated for 16 h before harvesting by centrifugation (15 min, 4000 g). The cell pellet was re-suspended in 30 mL lysis buffer (50 mM Tris-Cl pH 8.0, 500 mM NaCl, 10 mM imidazole, 0.5 mM TCEP) supplemented with complete protease inhibitor cocktail mini tablets (EDTA free; Roche), transferred to 50 mL falcon tubes and stored at –20°C.

Cell pellets were lysed using a cell disruptor (Constant Cell Disruption Systems) at 124 kPa, and the lysate was clarified by centrifugation at 30,000 ×g for 20 min at 4°C. The soluble portion was passed through a 5 mL IMAC column (HiTrap) pre-equilibrated with lysis buffer and bound protein washed with wash buffer (50 mM Tris-Cl pH 8.0, 300 mM NaCl, 20 mM imidazole). The recombinant protein was eluted in a linear gradient with elution buffer (50 mM Tris.Cl pH 8.0, 300 mM NaCl, 500 mM imidazole) and analysed by SDS-PAGE to estimate purity.

The pure recombinant protein was pooled, and the buffer was exchanged into PBS. Briefly, the protein sample was concentrated to 1 mL in a 10-kDa-molecular-weight-cutoff (MWCO) protein concentrator (VivaScience), resuspended with PBS buffer and reconcentrated. This was repeated twice to replace the elution buffer with PBS. Final protein concentrations were determined by measuring the absorbance of the samples at a wavelength of 280 nm using a NanoDrop spectrophotometer (Thermo Fisher Scientific, Waltham, MA, USA).

### Enzyme kinetics

An enzyme kinetics experiment was conducted using the absorbance method published by Zhong-Johnson*, et al.* [42]. In brief, 5 mg of PET powder and films were incubated in 2.5 ml of PBS buffer with varying enzyme concentrations: 50 nM, 100 nM, 150 nM, and 200 nM. The samples were incubated for 5 days on a shaking rack at 100 rpm and a temperature of 30 °C. Afterwards, 1.5 µl from each sample was assessed using a NanoDrop at a wavelength of 245 nm. This specific wavelength was chosen based on the absorption characteristics of mono-(2-hydroxyethyl) terephthalic acid (MHET) and terephthalic acid (TPA). Control samples, incubated with PBS buffer and PET but without enzyme, served as blanks for the NanoDrop readings. All measurements were taken in triplicate.

### Genome analysis

Before assembly, raw read quality was assessed using FastQC/0.11.7 [43]. Adapters were removed using Trimmomatic /0.39 [44] with the options “PE” and “-baseout trimmomatic ILLUMINACLIP:1_trimmomatic/adapt.fa:2:30:10”. A fasta file, “adapt.fa”, containing two sequences, was supplied as input to Trimmomatic to remove sequences from the TruSeq Universal adapter (AATGATACGGCGACCACCGAGATCTACACTCTTTCCCTACACGACGCTCTTCCG ATCT) and TruSeq Adapter Index 1(GATCGGAAGAGCACACGTCTGAACTCCAGTCACATCACGATCTCGTATGCCGT CTTCTGCTTG). Low-quality bases were trimmed using ‘Sickle’ [45] with the options “pe –-qual-type sanger –l 70”.

The high-quality reads were assembled using SPAdes/3.13.1 [46] with a k-mer range of 21 to 127. To assess assembly completeness, we used BUSCO/3.0.2 [47] with the Pezizomycotina dataset as a reference. The resulting assembly was further evaluated using QUAST/5.0.2 [48] to determine N50, L50, and genome size metrics.

To improve the accuracy of the assembly, we used BWA/0.7.17 [49] to align the raw reads to the assembly, followed by Pilon/1.23 [50] to perform error correction. The haplotigs in the assembly were identified and removed using ‘Purge Haplotigs’ [51]. Repetitive sequences in the genome were masked using ‘dustmasker’ [52] to reduce their impact on gene prediction. Finally, we performed gene prediction using AUGUSTUS/3.3.2 with default parameters [53]. The resulting genome assembly was used for further studies on this fungus.

The genome was annotated against several databases using BLAST [54] and HMMER3 [55], including KEGG [56], UniProt [57], Pfam [58] and TIGRfam [59]. Signal peptides were annotated using SignalP 5.0 software [60]. Additionally, all genome sequences were annotated using the InterProScan [61] tool with the –goterms flag, retrieving GO terms for each predicted protein sequence to investigate the biological significance of proteins of interest.

### RNA-Seq analysis

The quality of the raw RNA-seq reads was checked using FastQC/0.11.9 [43] and MultiQC/1.13 [62]. The raw reads were trimmed and filtered using fastp/0.23.2 [63] to remove low-quality bases and adapters. All parameters were set to their default value, but the following: “--trim_front1 9 –-trim_front2 9.”

The trimmed reads were aligned to the reference genome using Bowtie 2/2.4.5 [64]. The alignment file was sorted and indexed using SAMtools/1.10 [65]. The featureCounts program (Subread/1.5.3) [66] was used to count the number of reads mapped to each gene. The count table generated by featureCounts was imported into the R/4.2.1 [67] environment for statistical and differential gene expression analysis. The DESeq2 package [68] was used to identify differentially expressed genes. The R-based topGO package was used to perform gene Ontology (GO) enrichment analysis [69].

Transcripts with putative activity against PCL were identified by submitting the whole transcriptome to the webserver of the PlasticDB database [70], accessed on December 2024. The default values were used for the similarity search using the DIAMOND software version v2.0.6.144 [71] with an e-value cutoff of 1e^-6^ and 30% identity on the amino acid level. AlphaFold2 [72] was used to predict protein structures through the ColabFold [73] platform with default values. Predicted structures were annotated based on their structure similarity against all the data deposited in the PDB database [74] using the web service PDBeFold [75].

### Statistical analysis

Statistical analyses were performed using R Statistical Software version 4.3.1 [67] and Python version 3.10.12 [76]. A multivariate analysis assessed the impact of different experimental conditions on overall gene expression patterns. Principal Component Analysis (PCA) was conducted using the ‘prcomp’ function in R, focusing on the top 500 most variable genes to retain maximum variance. Subsequently, we employed the ‘adonis2’ function from the vegan version 2.6-4 [77] package in R to conduct a Permutational Multivariate Analysis of Variance (PERMANOVA). The analysis utilised a distance matrix based on PCA scores with the default Euclidean distance metric. A total of 999 permutations were executed to evaluate the significance of observed differences.

Statistical analyses were performed on two distinct datasets. The first dataset assessed differences in gene expression values among the three treatment groups: GLU, PCL, and GLUPCL. The second dataset investigated differences in clear zone areas among media types with various substrates. Initially, a one-way Analysis of Variance (ANOVA) was performed to assess overall differences among the treatment groups. Subsequently, Tukey’s Honestly Significant Difference (HSD) test was applied as a post hoc analysis to identify specific treatment pairs exhibiting statistically significant mean differences. The ANOVA and Tukey’s HSD test were conducted using Python with the scipy.stats (version 1.10.1; Virtanen*, et al.* [78]) and statsmodels (version 0.14.0; Seabold and Perktold [79]) libraries, respectively. The chosen significance level was α = 0.05 for both tests.

## RESULTS AND DISCUSSION

### Plastic degradation screening

The plastic-degrading abilities of *C. rosea* were investigated for four different plastic types: PCL, PLA, PS, and PE, emulsified in M9 minimal solid media. PET was not used due to its insolubility in all evaluated solvents. After two days of incubation, *C. rosea* exhibited a clear degradation halo on PCL-emulsified agar plates, indicating its ability to degrade PCL. However, there was no visible evidence of degradation on PLA, PS, and PE-emulsified agar plates after 21 days. No microorganism has yet been reported capable of degrading all four plastic types tested here. However, by searching the PlasticDB database, it is possible to find 36 microorganisms out of 633 in the database that can degrade both PCL and PLA [70].

Following the identification of the PCL-degrading abilities of C. rosea, we investigated whether glucose supplementation affected PCL degradation. In biodegradation assays conducted over 18 days (**Figure 1**), the M9 medium alone had a mean degradation area of 1048 mm² (± 53 StDev), demonstrating that C. rosea could grow using PCL as a sole carbon source. C. rosea did not grow on an agar-only medium, indicating it cannot use agar as a carbon source. Notably, PCL biodegradation was completely inhibited when glucose was added to the growth medium, with no evidence of biodegradation observed over 18 days. Additional substrate screening experiments examining the effects of various carbon sources on PCL degradation indicated that C. rosea with PCL-only medium may be carbon starved (**Figure S1**).

**Figure 1:**
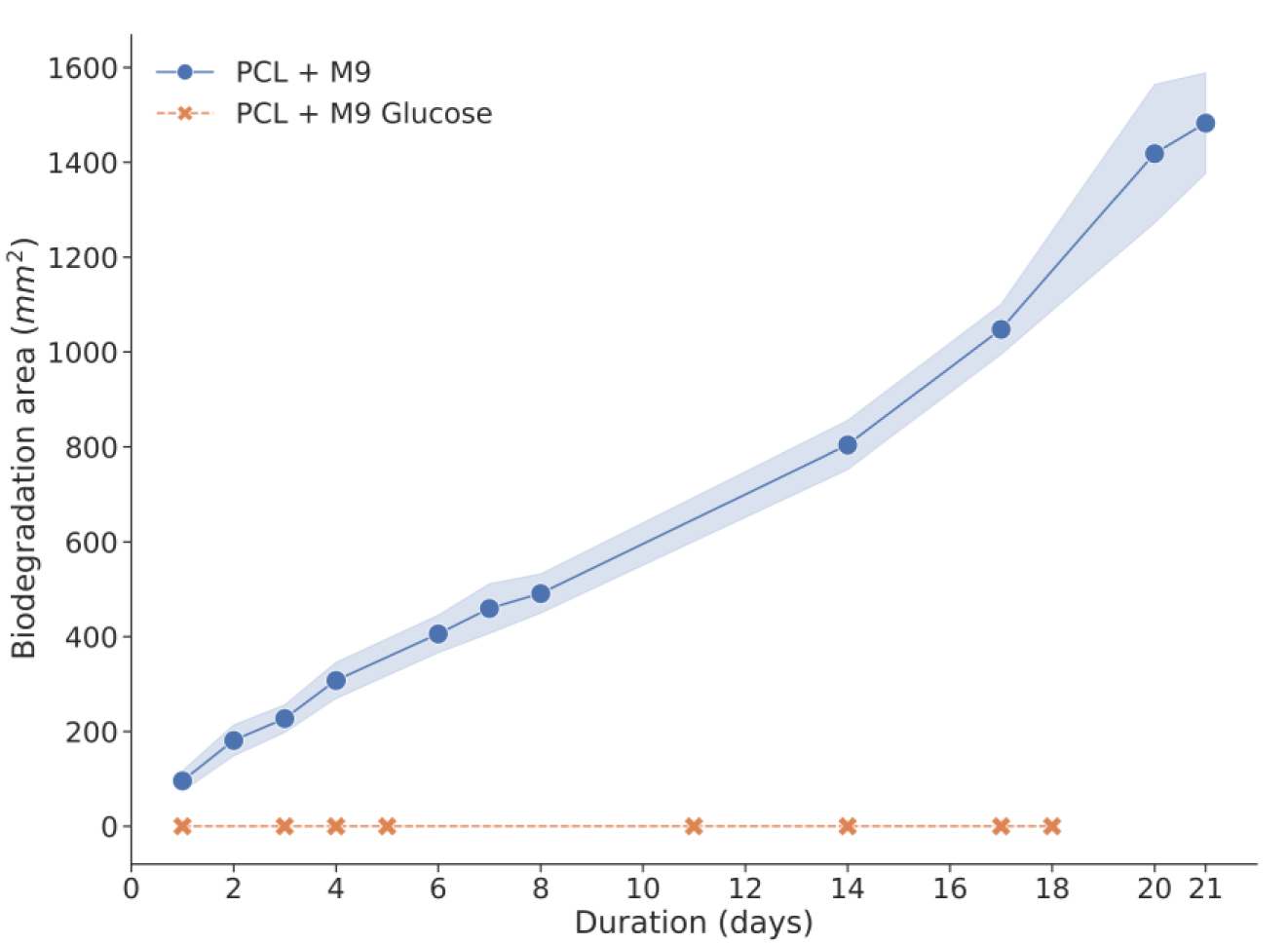
Ability of Clonostachys rosea to degrade a PCL emulsion in M9 media with and without glucose supplementation. Biodegradation was quantified by measuring the area of the clear halo formed around the microbial colonies. Each treatment was performed in triplicate. Shaded error bands represent standard deviation values.

The inhibition of PCL degradation in the presence of glucose may be attributed to glucose being a preferred and readily metabolisable carbon source compared with PCL. When glucose is available, the fungus may not produce the necessary enzymes for PCL degradation, leading to the complete inhibition of PCL degradation. This carbon catabolite repression is a common phenomenon and has also been reported in the literature for other carbon sources besides glucose [82–84].

It is not uncommon to find examples in the literature where the plastic biodegradation of microorganisms is overstated [85], with microbes only degrading a small percentage of the plastic or degrading synthetically aged plastics. Therefore, we confirmed the degradation of solid PCL material, rather than emulsions by *C. rosea,* by conducting experiments using PCL films. Further, we confirmed the impact of glucose on PCL film degradation by *C. rosea,* using three distinct liquid media: M9, M9 with yeast extract (as a standard growth supplement), and M9 with glucose. Yeast extract was chosen as a more conventional growth medium supplement to ensure consistent microbial development, as it is frequently shown to improve biodegradation rates [41]. After a 180-day incubation at 28°C, *C. rosea* completely degraded (100%) PCL films in the M9 with yeast extract medium. In the M9 medium alone, 59.25% (SD: 7.10%) of the PCL films were degraded. In contrast, exposure to the medium supplemented with glucose resulted in only a 1.89% (SD: 0.58%) weight loss of the PCL film (**Figure 2**).

**Figure 2:**
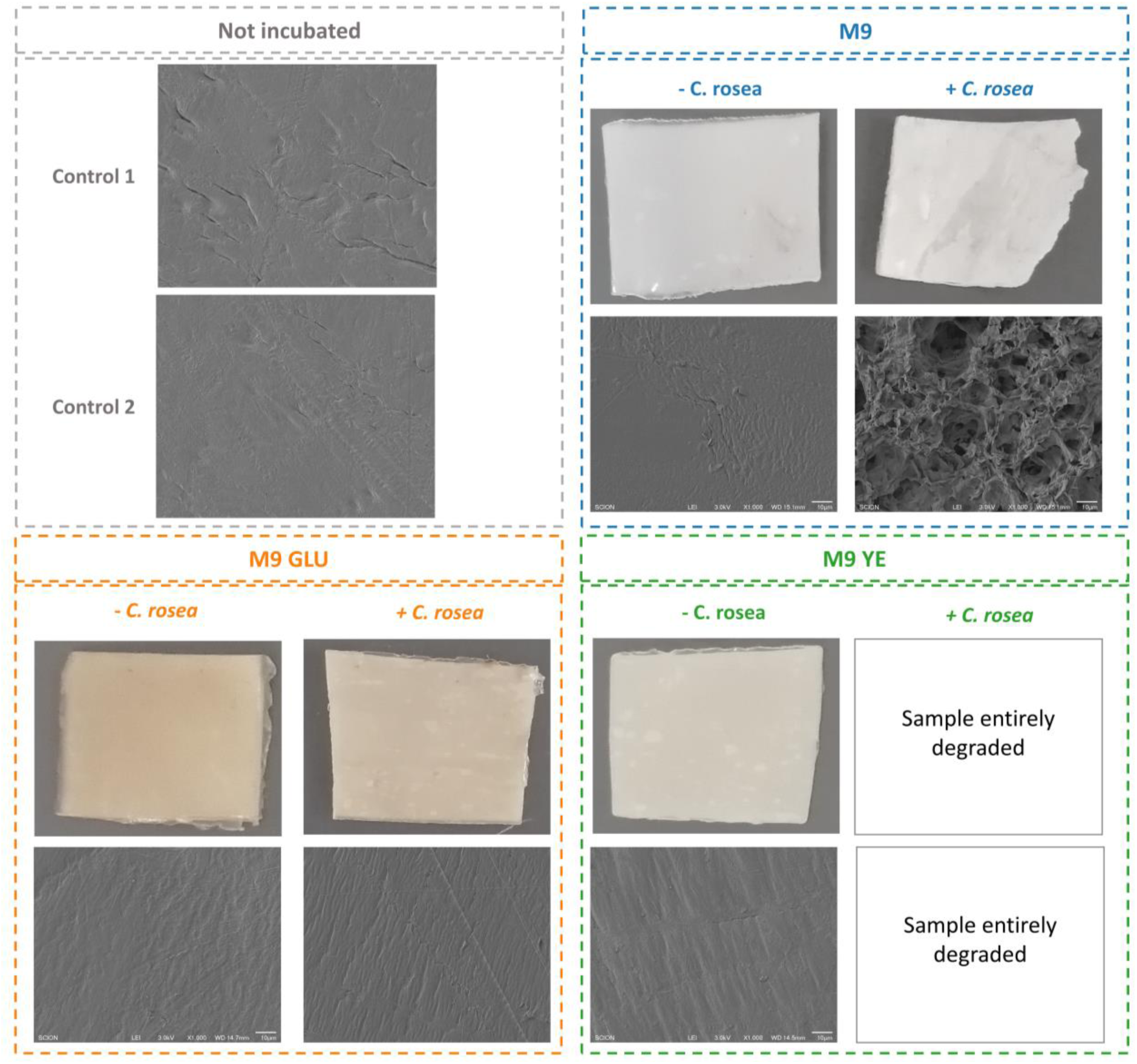
Biodegradation assay of PCL films by *Clonostachys rosea*. The fungus degraded PCL completely in the medium with yeast extract (YE), partially in the medium with M9 only, and did not degrade any PCL in the medium with glucose. The assay was conducted over a 180-day incubation period at 28 °C on M9 liquid media supplemented with different substrates: M9, M9 + yeast extract (4 g/L), and M9 + glucose (4 g/L), with all conditions replicated four times. The film weights were weighed before and after incubation to determine degradation percentages. Scanning electron microscopy (SEM) was performed on control and incubated samples at 1000x magnification.

Whether incubated or not, scanning electron microscopy images confirmed that control samples (samples not incubated or incubated without *C. rosea*) had a smooth surface morphology. In the case of samples subjected to *C. rosea* incubation in the M9 glucose medium, no discernible differences were observed compared with the control samples, and signs of biodegradation were notably absent. In contrast, samples incubated with *C. rosea* on the M9 medium alone showcased a roughened surface characterised by cracks and voids. Samples incubated with *C. rosea* on the M9 yeast extract medium underwent complete degradation, precluding their inclusion in the SEM analysis.

### Genome analysis

To understand the genetic mechanisms involved in the ability of *C. rosea* to degrade PCL, we sequenced the fungus’ genome. The sequencing of the *C. rosea* genome resulted in 8,524,227 paired reads (**Table S1**). After quality control steps to remove low-quality reads, 98.75% of reads were retained. This retention rate suggests that reads used in consecutive analyses were of high quality. The size of *C. rosea’s* assembled genome is 56.8 Mbp, with 351 contigs and an average coverage of 31x (**Table S2** and **Figure S2)**. This estimation aligns well with previous reports of the *C. rosea* genome size, which ranges from 55.4 to 58.3 Mbp [36, 86]. The N50 metric of 378.5 Kbp compares broadly with previously published values for *C. rosea* genomes of 569 Kbp [36] and 790 Kbp [37], and indicates a highly contiguous assembly. Furthermore, we analysed the assembled genome to assess its content with respect to universal single-copy orthologs from the Pezizomycotina subphylum. This analysis verified that the assembled genome encompassed a significant portion of the anticipated universal single-copy orthologs. Specifically, 3116 of 3156 orthologs were identified as present and complete (**Figure 3A**). DNA extraction from fungal mycelia targeted the haploid life stage of *C. rosea* [87]This was evident in the BUSCO results, which showed abundant single-copy genes (3092 genes) and few duplicates (24 genes). The BUSCO and QUAST analysis results indicated that genome assembly did not contain any contamination, which was also confirmed by the NCBI FCS (Foreign Contamination Screen) pipeline [88].

**Figure 3:**
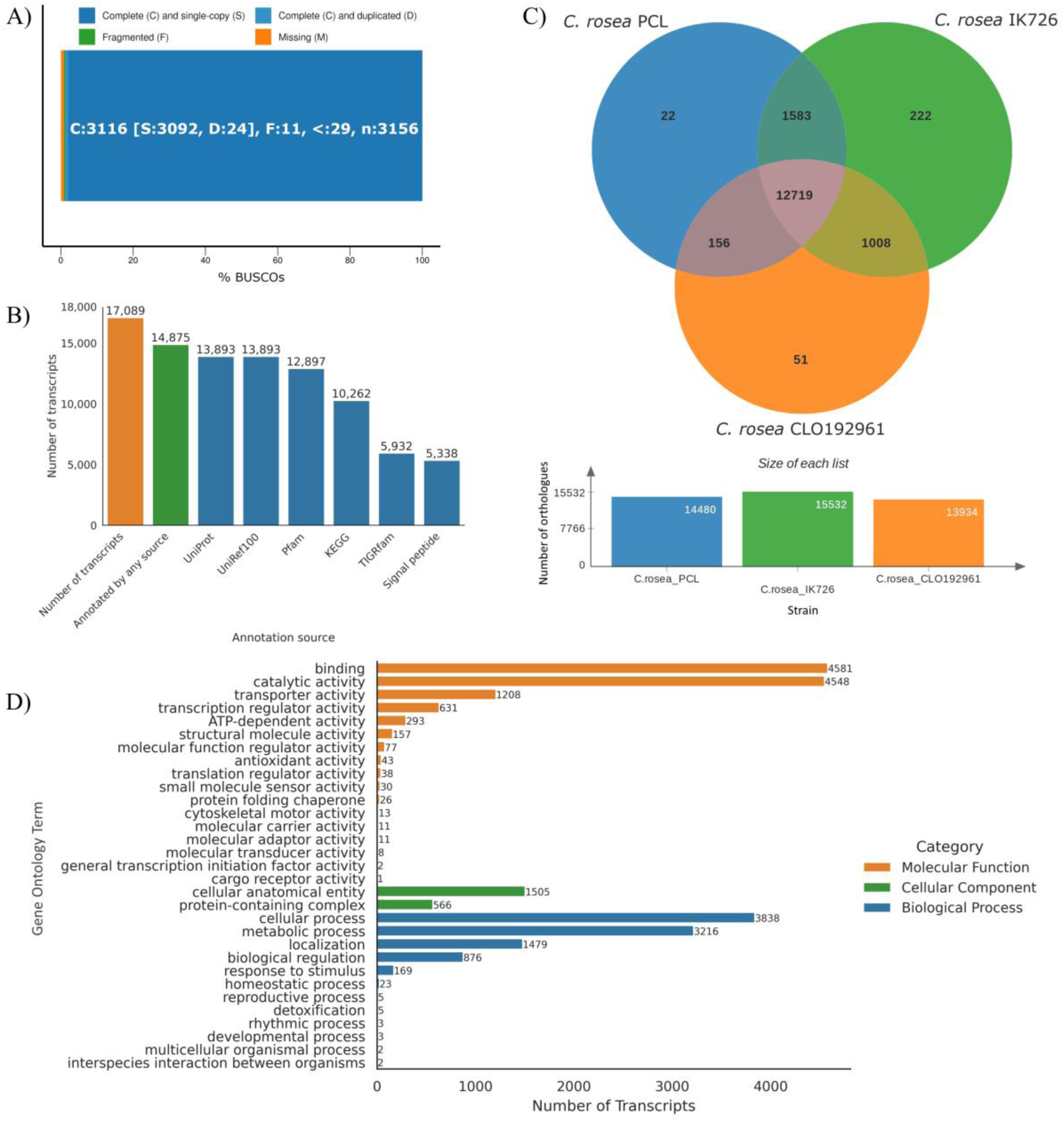
A) BUSCO report of genome assembly completeness. B) Number of annotated transcripts in the *C. rosea* genome for each annotation source. C) Orthologous clusters shared between the sequenced genome from this study (*C. rosea* PCL) and two published *C. rosea* genomes. D) Distribution of gene ontology (GO) terms annotated on the *C. rosea* genome.

A total of 17,089 genes were predicted in the genome of *C. rosea.* Overall, 2,214 genes (13%) were unannotated, with 14,875 genes (87%) annotated across all annotation sources (**Figure 3B**). The annotation revealed 5,338 signal peptides involved in directing proteins to the correct location in the cell through translocation [89]. Annotation with the UniProt and UniRef100 databases gave the highest number of annotations (13,893 each). The Gene Ontology annotation of the *C. rosea* genome, which provides insights into the biological processes, molecular functions, and cellular components, suggests that, as most genomes [90], a significant portion of genes encode fundamental cellular activities like metabolic and cellular processes, as well as the spatial arrangement within the cell (**Figure 3D**).

To compare the genome of the *C. rosea* strain described here (hereafter referred to as *C. rosea* PCL) to other published *C. rosea* genomes, i.e., strains IK726 and CLO192961 (NCBI Genome Assembly accession numbers GCA_902827195.2 and GCA_902085965.1, respectively), we used the OrthoVenn3 tool (**Figure 3C**). Out of the total clusters of orthologues identified in each strain, there were 14,480 in strain PCL, 15,532 in IK726, and 13,934 in CLO192961. In addition, 12,719 clusters were shared among all three strains, emphasising these as core genomic elements underlying the biology of these strains. Interstrain comparative analysis revealed varying levels of shared orthologues between pairs of strains. There were 1,583 shared clusters between PCL and IK726, 156 between PCL and CLO192961, and 1,008 between CLO192961 and IK726. These pair-wise shared clusters indicate that strain PCL is more similar to strain IK726, making strain CLO192961 the most distinct among the analysed genomes. Each strain also had unique orthologue clusters: PCL had 22 unique clusters, IK726 had 222, and CLO192961 had 51, illustrating genetic diversity among the strains. However, it’s crucial to note that if the genomes are incomplete, then this observed diversity could be influenced by the incompleteness of the genomes.

### Gene expression profiling

Next, we conducted an RNA-Seq experiment to identify putative genes and pathways for PCL degradation in C. rosea and the mechanisms for glucose-mediated inhibition of this process (**Figure 4A**). We cultivated *C. rosea* for six days on the following solid media: M9 with PCL, M9 with glucose, and M9 with PCL and glucose. Next, RNA was extracted (mean RIN values of 7.04, SD: 1.46), and expression analysis was conducted. A total of 16,807 genes had mapped reads. No genes were notably downregulated in the GLUPCL medium (M9 medium with both glucose and emulsified PCL) compared with GLU medium control (M9 medium and glucose), while two genes were upregulated. Thus, adding emulsified PCL to the glucose medium did not significantly affect the gene expression profile of the fungus. In contrast, there were 1835 significantly downregulated genes and 2184 upregulated genes in the PCL medium (M9 medium with emulsified PCL) compared with the GLUPCL medium. Similarly, the comparison between PCL and GLU media showed 1783 significantly downregulated genes and 2228 upregulated genes in the PCL media (**Figure 4B**), showing that the two treatments with glucose generated a very different gene expression profile compared to the PCL-only treatment.

**Figure 4:**
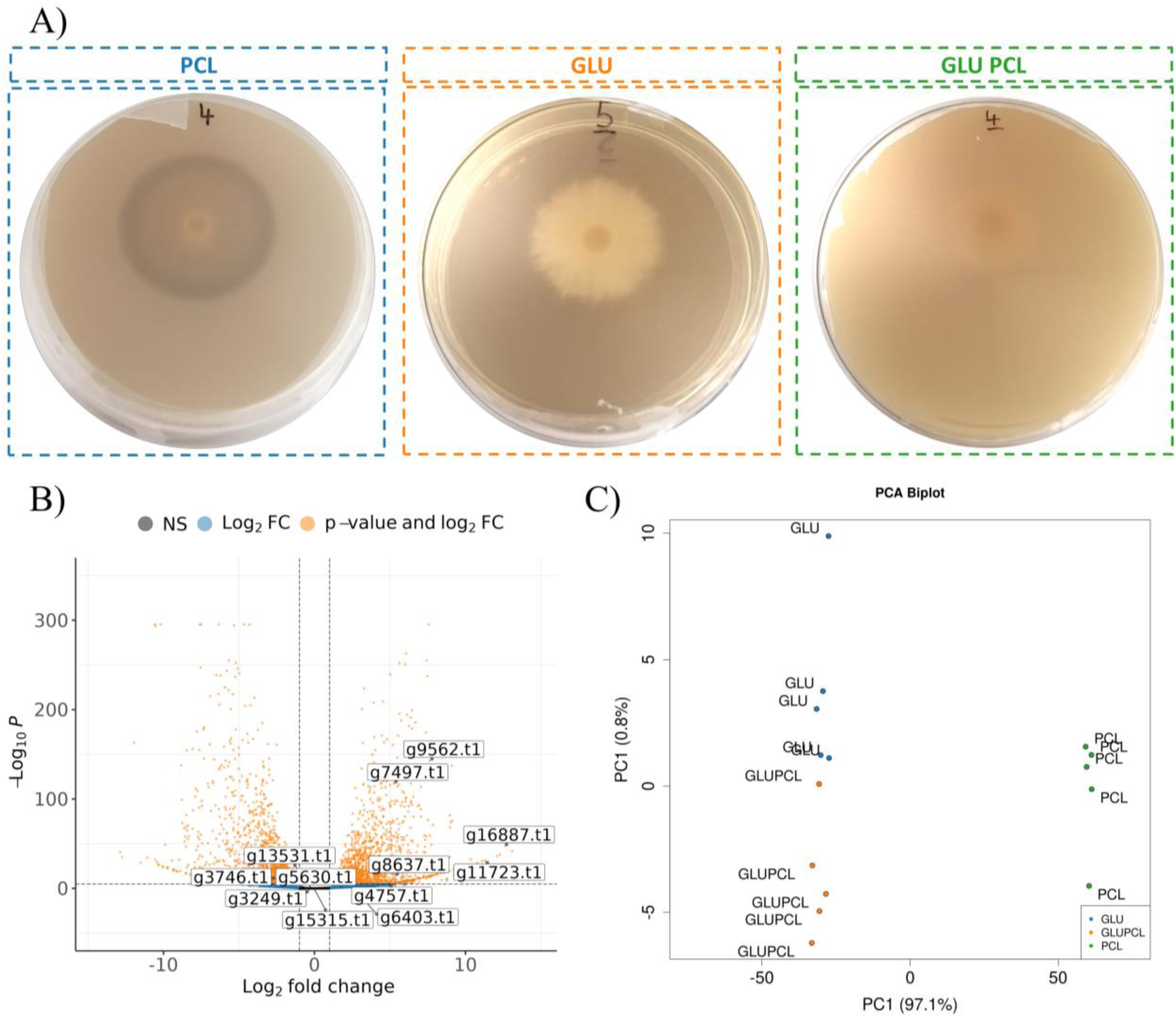
A) *Clonostachys rosea* growing on the three media types used for the RNA-Seq experiment: (i) M9 agar with polycaprolactone (PCL), (ii) M9 agar with glucose (GLU), and (iii) M9 agar with both glucose and PCL (GLUPCL). All treatments had five replicates (n = 15). Clear zones can be observed on PCL-only plates but not on plates containing glucose. B) Differentially expressed transcripts when comparing PCL versus GLU treatments, with cutoffs of FDR < 0.1 and absolute Log_2_FoldChange > 1. Positive Log_2_FoldChange values indicate the transcripts that were upregulated by the PCL treatment relative to the GLU treatment. Genes with labels were annotated as putatively encoding for PCL-degrading enzymes according to the PlasticDB database. C) Principal Component Analysis (PCA) biplot of the expression values for the top 500 most variable genes. Significant differences in gene expression patterns among treatments were observed (R² = 0.976, p < 0.001) based on PERMANOVA with 999 permutations using the Euclidean distance metric.

A Principal Component Analysis (PCA) Biplot showed that samples from the three treatments had compositionally distinct gene expression profiles (**Figure 4C**). Samples with and without glucose separated principally across the x-axis, which accounts for a substantial 97.1% of the total data variance, further illustrating the influence of glucose on *C. rosea* gene expression. While GLU and GLUPCL samples clustered together on the x-axis, they separated along the y-axis. However, the y-axis explained only 0.8% of the total data variance, implying that the difference caused by PCL presence was small compared to the effect of glucose addition.

As discussed earlier, *C. rosea* degradation of PCL was inhibited in the presence of glucose. Thus, biological processes upregulated in the PCL treatment compared with the GLU treatment may better elucidate the potential degradation pathways of PCL (**Figure 5A**). Therefore, we performed a Gene Ontology enrichment analysis and found the upregulation of several biological processes, including carbohydrate metabolic process, fucose metabolic process, and xylan catabolic process in the presence of PCL and absence of glucose. We also found upregulation of genes involved in aromatic amino acid metabolism, such as L-phenylalanine and tyrosine, and catechol-containing compound metabolic process. Those upregulated biological processes may indicate that *C. rosea* may mobilise pathways to metabolise the aromatic components produced by the breakdown of the PCL polymer, such as ε-caprolactone. We also found several downregulated biological processes during PCL degradation, including polysaccharide and inositol catabolic processes, proteolysis and fatty acid biosynthetic processes (**Figure 5B)**.

**Figure 5:**
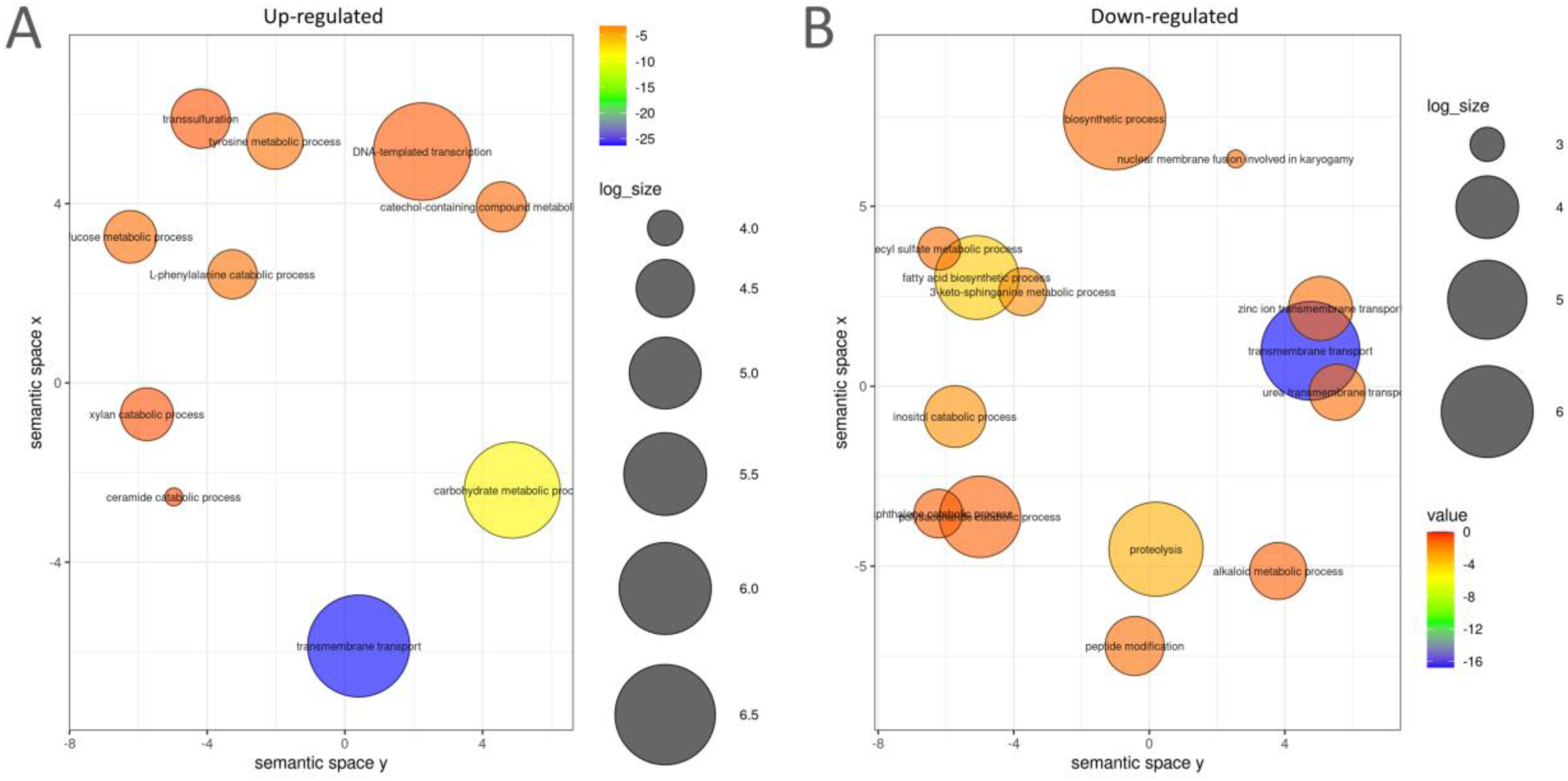
Gene Ontology (GO) enrichment analyses summarised and visualised using REVIGO. Significantly enriched GO terms related to biological processes: A) up-regulated and B) down-regulated transcripts comparing PCL versus GLU-GLUPCL. GO terms are represented by circles and are clustered according to semantic similarities to other GO terms. Circle size is proportional to the frequency of the GO term, whereas colour indicates the log10 P-value for the enrichment derived from the topGO analysis.

The annotation of PCL-upregulated genes using the KEGG database reinforces insights gleaned from the Gene Ontology analysis, identifying numerous genes involved in the fatty acid degradation pathway (**Figure 6**). Fatty acid degradation, or beta-oxidation, involves breaking down long-chain fatty acids into acetyl-CoA [91]. This versatile metabolite feeds into the tricarboxylic acid (TCA) cycle, generating ATP and reducing equivalents [92]. The upregulation of genes within this pathway in the PCL treatment suggests that *C. rosea* may exploit similar enzymatic machinery to metabolise the products generated during PCL degradation. This pathway has also been suggested to play a role in plastic degradation by other fungi [93]. Simultaneously, the presence of genes from the fatty acid degradation pathway could indicate a metabolic adaptation for energy production under the PCL conditions. In the previous Gene Ontology enrichment analysis, fatty acid biosynthetic processes were downregulated, suggesting that the degraded fatty acids likely originate from PCL degradation rather than produced by *C. rosea*. Fungi typically accumulate substantial quantities of lipids within both their hyphae and spores. These lipids are crucial as carbon and energy reservoirs, particularly during periods of starvation and spore germination [94–96]. Therefore, with the potentially reduced availability of traditional energy sources such as polysaccharides, as suggested by the downregulation of polysaccharide catabolic processes in the GO analysis, *C. rosea* might try to compensate by upregulating fatty acid degradation of internal resources or PCL degradation products.

**Figure 6:**
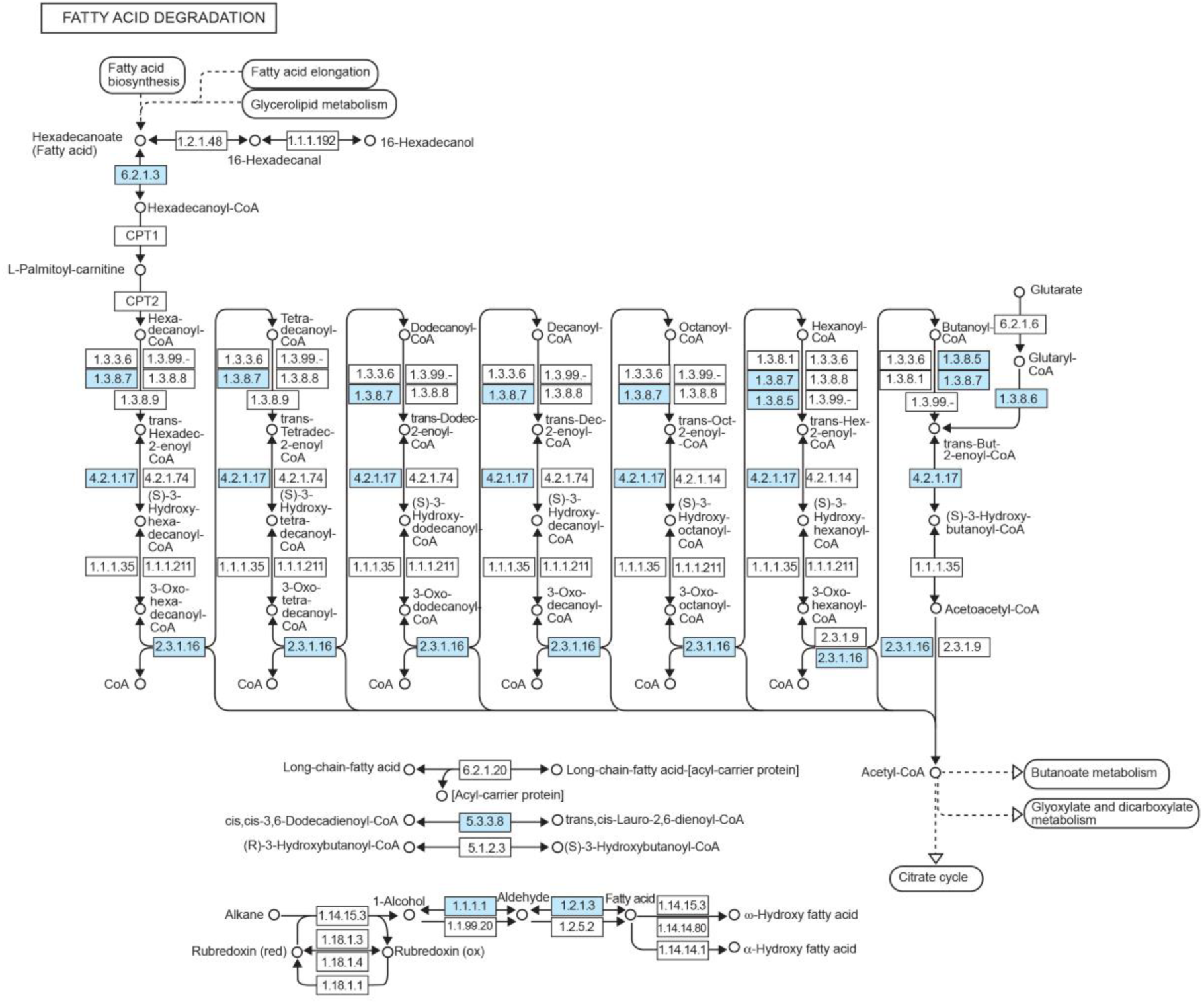
Upregulated expressed genes mapping to the fatty acid degradation pathway. Blue rectangles represent reactions upregulated in PCL compared to GLU and GLUPCL treatments, with cutoffs of FDR < 0.1 and absolute Log_2_FoldChange > 1.

### Heterologous expression and enzyme kinetics

We identified 12 *C. rosea* genes as putatively coding for PCL-degrading enzymes by annotating all transcribed genes against PlasticDB, a database for genes reported to be involved in plastic biodegradation [70]. Two transcribed genes, g16887.t1 and g9562.t1, stood out for being significantly differentially expressed and having high expression levels in the presence of PCL, while having an absence of expression in glucose treatments (**Figure 7**; g16887.t1 log2FoldChange=13.14; g9562 log2FoldChange=11.96). These genes were, therefore, considered key candidates for PCL degradation. We also searched for these genes within the genomes of the *C. rosea* strains IK726 and CLO192961 using BLAST [54] with a cutoff e-value of 1e-6 and a minimum per cent identity of 30% at the amino acid level. Strain IK726 contained all 12 genes in its genome, while strain CLO192961 possessed 10.

**Figure 7:**
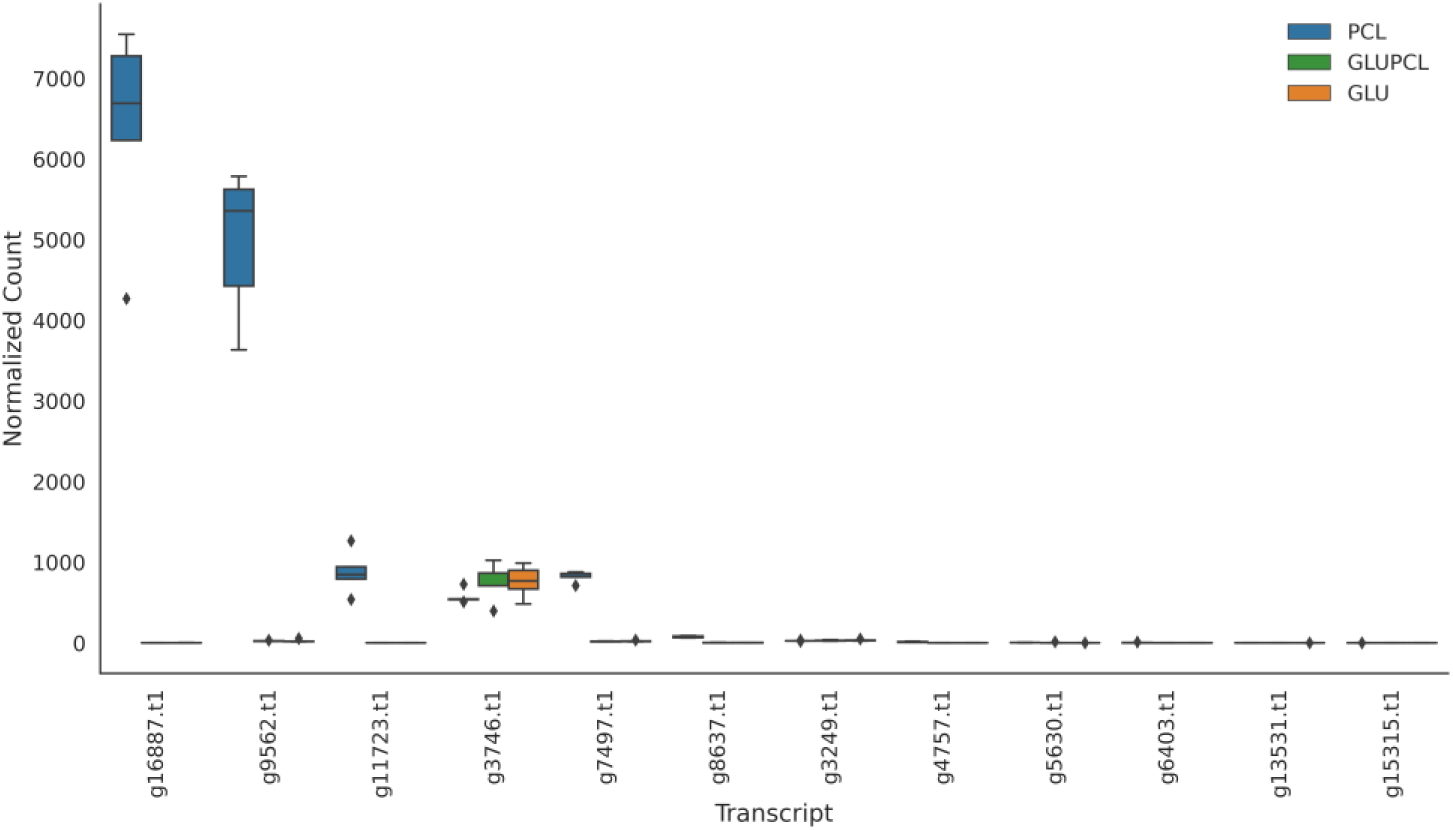
Expression of *Clonostachys rosea* transcripts annotated with putative PCL degradation activity by the PlasticDB tool. ANOVA (F = 14.96, p < 0.0001) indicated overall differences. Tukey’s HSD revealed significant differences between GLU and PCL, and GLUPCL and PCL treatments, but not between GLU and GLUPCL. The box represents the range between the 25th percentile and the 75th percentile of the data. The line inside the box represents the median. The whiskers indicate the range of the data that are not considered outliers. Individual data points that fall outside the whiskers are outliers.

The expressed g9562.t1 gene was identified by InterProScan as an alpha/beta-hydrolase. Based on PlasticDB, it was 66.8% identical to a cutinase from *Humicola insolens* (PlasticDB code: PLDB00121), demonstrated to degrade PET [97]. Similarly, the expressed gene g16887.t1 was annotated by InterProScan as an alpha/beta-hydrolase. Based on PlasticDB, it was 68.3% identical to a cutinase from *Thermobifida fusca* (PlasticDB code: PLDB00073), another fungus reported to degrade PET [98]. The action of cutinases, esterases and hydrolases in plastic degradation is well documented, particularly due to their ability to hydrolyse ester bonds, a key structural feature of many plastics, including PCL [99, 100].

To confirm the ability of the enzymes coded by these two expressed genes to degrade PCL, we expressed the genes for these proteins in Shuffle T7 Competent *E. coli* cells and purified the resultant recombinant proteins via affinity chromatography. The molecular weight (Mw) for recombinant g9562.t1 was 20.0 kDa and 34.1 kDa for g16887.t1. An aliquot (1 µL at a concentration of 2 µg/µL) of the purified enzymes was applied onto PCL-emulsified agar for 24 h at room temperature (18-24 °C) and clear zones were observed for both enzymes, confirming their activity against PCL. The control with ddH_2_O showed no signs of degradation (**Figure S7**).

Various studies report that some enzymes capable of degrading PCL also possess the ability to degrade PET [23, 81, 101]. The ability of these enzymes to degrade PET was therefore tested by an absorbance method [42]. Only the enzyme expressed by the transcribed gene g9562.t1 exhibited concentration-dependent catalytic activity against PET, with higher enzyme concentrations leading to increased substrate degradation, effectively breaking down PET film and powder substrates over a 5-day incubation period (**Figure 8C**). The enzymes coded by the transcribed genes g9562.t1 and g16887.t1 were deposited in the PlasticDB database as cutinases. g9562.t1 (accession: PLDB00220) was deposited with PCL and PET degradation activity, while g16887.t1 (accession: PLDB00221) was deposited with PCL degradation activity only.

**Figure 8:**
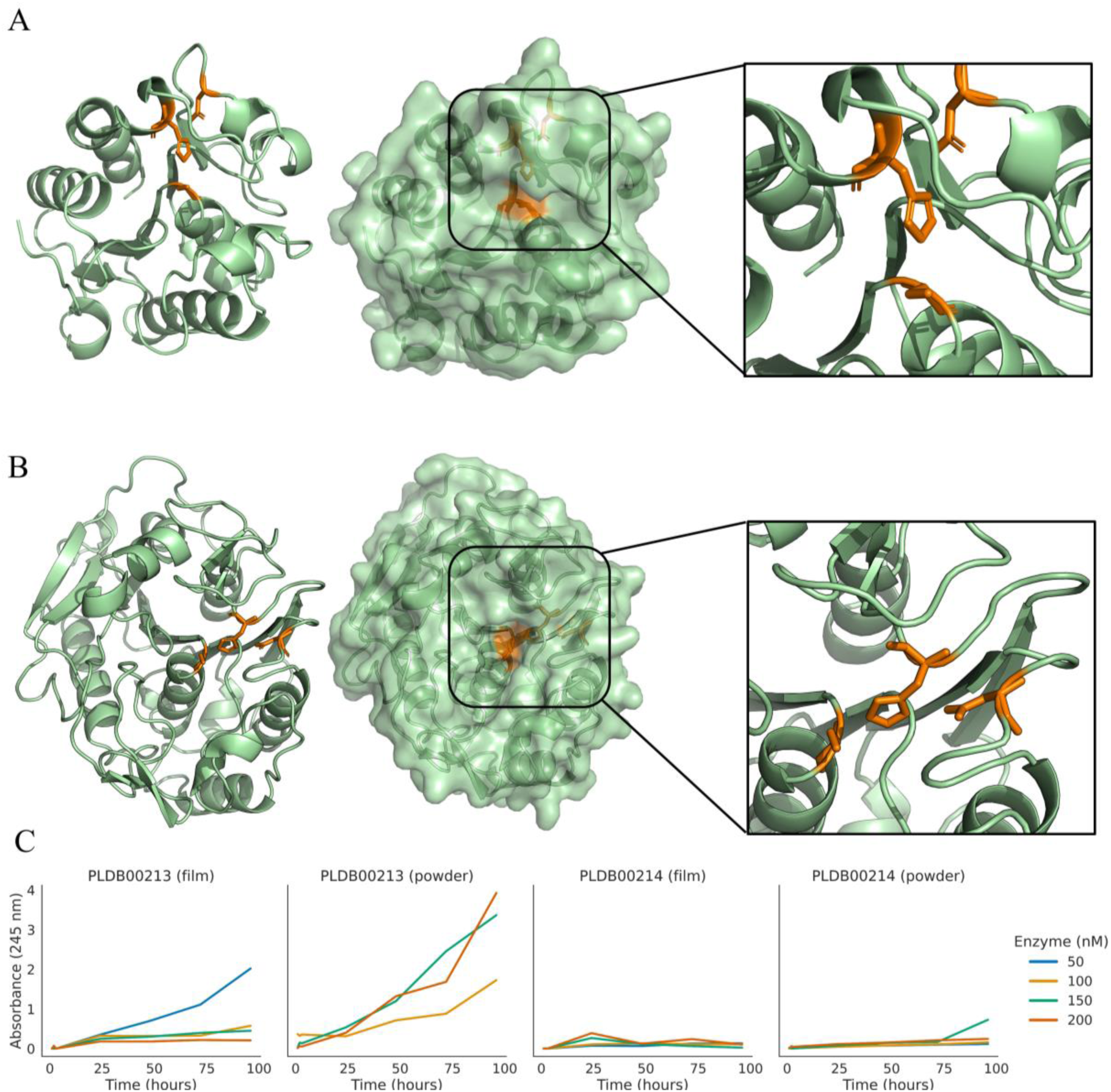
Alphafold2 prediction of protein structures. A) The cutinase PLDB00220 (g9562.t1) showed activity against PCL and PET. AlphaFold2 prediction scores: pTM =0.85 and pLDDT=91.4). B) The putative cutinase PLDB00221 (g16887.t1) showed activity against PCL only. AlphaFold2 prediction scores: pTM=0.92 and pLDDT=94.9). C) enzymatic degradation kinetics of PET powder and film using both purified enzymes. Degradation rates were determined using absorbance at representative enzyme concentrations; absorbance values are the mean of three readings per sample. Higher absorbance values indicate higher enzymatic activity against PET.

We used the algorithm AlphaFold2 to predict the structure of enzymes PLDB00220 (g9562.t1) and PLDB00221 (g16887.t1). Using the predicted structure, we annotated both proteins using the PDBeFold web tool. PLDB00220 matched a cutinase from *H. insolens* (PDB: 4oyy; 65% sequence identity). Enzyme PLDB00220 active site is at residues SER 131, with HIS 199 and ASP 186 associated residues (**Figure 8A**). It is in a long cleft and likely to accommodate longer polymer substrates. A putative mobile loop near the cleft may further expand the space available for substrate binding. Numerous similar structures of different cutinases share a similar substrate binding groove, indicating a highly conserved functionality across species.

Enzyme PLDB00221 matched a polyhydroxybutyrate depolymerase from *Penicillium funiculosum* (PDB: 2d81; 69.2% identical) that was shown to degrade PHB and a trimer of R3HB [102]. The enzyme PLDB00221 active site is predicted to be at residues SER 39, with HIS 155 and ASP 121 associated residues (**Figure 8B**). It is situated in a more constrained pocket when compared to the other enzymes. This smaller pocket may explain why PLDB00221 was only active against PCL, not PET. Furthermore, PLDB00221 and its *P. funiculosum* homolog (PDB: 2d81) share nearly identical residues lining the substrate binding pocket/groove. This structural similarity suggests that both enzymes likely exhibit comparable catalytic activity against polyesters.

## CONCLUSION

We demonstrate *C. rosea’s* ability to biodegrade PCL and PET, highlighting the influence of substrate composition on the biodegradation process. The biodegradation assays revealed that the carbon sources present significantly impact the degradation process, with potato starch providing the most favourable conditions out of those tested. We demonstrated that the fungus underwent a significant shift in metabolic activity in the presence of PCL. Gene expression experiments highlighted the possible importance of the fatty-acid biodegradation pathway for plastic biodegradation. The analysis of upregulated transcripts against the PlasticDB database revealed two transcripts showing high expression levels in PCL conditions, suggesting their possible role in *C. rosea* PCL degradation phenotype. Both transcripts were heterologously expressed, and the produced enzymes exhibited activity against PCL, while one of the enzymes also demonstrated activity against PET polymers. Taken together, the knowledge generated in this study can aid in developing more efficient microbiological strategies for plastic degradation and recycling by optimising carbon sources, metabolic pathways, and enzyme activity on plastic substrates. Our findings emphasise the promising role of fungi in bioremediation and open doors to further research into enhancing the ability of fungi to degrade different types of plastics.

## DATA AVAILABILITY

The assembled genome, the raw genome Illumina paired-end sequences, and the raw RNA-Seq Illumina paired-end sequences have been submitted to the NCBI database, accession number PRJNA1017662, https://www.ncbi.nlm.nih.gov/sra/PRJNA1017662. The g9562.t1 and g16887.t1 protein sequences and their AlphaFold2 predictions were submitted to the PlasticDB database: https://plasticdb.org/proteins_00229 and https://plasticdb.org/proteins_00230, respectively. The code used for all bioinformatics and statistical analysis is organised in Jupyter notebooks deposited into Figshare: https://doi.org/10.17608/k6.auckland.24452110.

## ACKNOWLEDGEMENTS

V.G. was supported by a Ph.D. stipend from the George Mason Centre for the Natural Environment (New Zealand). Additional support was provided by the Aotearoa Impacts and Mitigation of Microplastics (AIM2) project (Ministry of Business, Innovation, and Employment, New Zealand, Endeavour Fund C03X1802) and the Engineering Microbial Enzymes for Plastic Recycling and Environmental Remediation project (Ministry of Business, Innovation, and Employment, New Zealand, Smart Ideas Fund, UOAX1916).

